# Evaluation of the efficacy of the protopine total alkaloids of *Macleaya cordata* (Willd.) R. Br. in controlling *E. coli* infection in broiler chickens

**DOI:** 10.1101/2024.07.03.601902

**Authors:** Zhen Dong, Yufeng Xu, Zhiqin Liu, Jianguo Zeng

## Abstract

**Objective:** This study was carried out to investigate the preliminary evaluation of the effectiveness of protopine total alkaloids of Macleaya cordata (Willd.) R. Br. (MPTA) extract in the control of artificially infected avian pathogenic E. coli in the peritoneal cavity of chickens.

**Methods:** In this test, Lingnan yellow hybrid chickens (male, 10 days old) were attacked with E. coli O78 and then treated orally with different concentrations (25 - 1600 mg/kg) of MPTA Pulvis (MPTA-P) and 0.5% Siweichuanxinlian Powder (SWCXL-P).

**Results:** The results showed that different concentrations of MPTA-P and SWCXL-P were effective in reducing the mortality of E. coli and promoting the recovery of the affected organs, with the best intervention being the supplementation of 400-1600 mg/kg of MPTA-P for 7 consecutive days. It has been concluded that the addition of 400 mg/kg MPTA-P for 7 days reduces the severity and mortality and accelerates the recovery process of E. coli disease in chickens and has a protective effect against organ lesions caused by E. coli infection.

**Limitations:** The study lacked comparisons of carrier populations and characterization of inflammatory markers.

**Conclusions:** MPTA may be a potential alternative drug for the prevention or treatment of avian E. coli disease.

## 1. Introduction

Avian pathogenic *Escherichia coli* (APEC) is one of the most common bacterial pathogens of poultry in the farming industry (Johnson *et al*., 2022). APEC infections can be characterized by local and systemic infectious inflammatory conditions such as salpingitis, pneumonia, arthritis, peritonitis, septicemia and respiratory infections (Dho-Moulin and Fairbrother, 1999). APEC can be a threat to humans through the food chain via contaminated poultry products, water and soil (Jørgensen *et al*., 2022; Rabiu *et al*., 2022; Sarowska *et al*., 2022). APEC strains are genetically similar to some human-derived *E. coli* and can cause digestive tract infections, reproductive tract infections and meningitis in rodents (Meena *et al*., 2021; Sarowska *et al*., 2022). Poultry of all ages are susceptible hosts for APEC, and APEC infections can be found in almost all types of production systems in chickens, turkeys, ducks and geese (with the highest infection rate of over 30% in laying hens) (Dube and Mbanga, 2018; Lutful Kabir, 2010), resulting in suppressed body weight gain and reduced feed conversion, which can be severe and can be life-threatening and cause significant economic losses (Helmy *et al*., 2022). Thus, APEC poses a major threat to global poultry production.

Modern farming is usually treated with antibiotics and vaccines against APEC infections (Christensen, Bachmeier, and Bisgaard, 2021). The overuse of antibiotics and the use of antibiotic growth promoters has led to the emergence of a large number of resistant strains of bacteria, thus limiting the effectiveness of antibiotic treatment (Dos Anjos Adur *et al*., 2022; Oliveira *et al*., 2022). The diversity of bacterial strains and cross-contamination with heterologous serotypes has also led to a large number of vaccine failures (Ghunaim, Abu-Madi, and Kariyawasam, 2014; Koutsianos *et al*., 2022). In addition, APEC is an important zoonotic food-borne pathogen. Therefore, the development of new and effective methods for the control of APEC infections in poultry is of great public health relevance.

Novel therapeutic approaches, such as probiotic supplements and phage and quorum sensing inhibitors, are being actively developed. (Eid *et al*., 2022; Guo *et al*., 2014; Helmy *et al*., 2018; Liang *et al*., 2021). Drawing on traditional ethnomedicine and the development of effective therapeutic medicines from natural plants is one of the current possibilities. Standardized plant extracts or purified active compounds (preparations) can be used as alternatives to antibiotics after they have been evaluated for safety and efficacy (Dakheel *et al*., 2020; Kulnanan *et al*., 2021; Zhong *et al*., 2014). *Macleaya cordata* (Willd.) R. Br. (*M. cordata*) is a perennial plant of the opium poppy family mainly found in China. The fruit pods and roots are the main parts used in traditional medicine. The greatest pharmacological activity in *M. cordata* comes from alkaloids, of which four are the most abundant, namely, sanguinarine (SAN), chelerythrine (CHE), protopine (PRO) and allocryptopine (ALL). *M. cordata* extract (MCE), of which SAN and CHE are the main components, has been widely used in livestock production and agricultural pest control (Ke *et al*., 2017, 2019; Zhang *et al*., 2021). Recently, significant anti-inflammatory and antioxidant activity has been reported for the protopine alkaloid components (PRO and ALL) of this plant (Bae *et al*., 2012; Dong *et al*., 2022; Muñoz *et al*., 2011). In addition, the addition of Bopu Powder®, which is based on the protopine total alkaloids of *M. cordata* (MPTA), to feed can improve egg quality, antioxidant capacity and intestinal health in hens (Liu *et al*., 2022). Previously, oral administration of MPTA significantly reduced diarrhea in the treatment of *E. coli*-induced infectious enteritis in mice (unpublished). However, it is not clear whether MPTA is effective in controlling APEC infection in poultry. Based on the above facts, the present study reports on the control effect and effective dose screening of MPTA in broiler chickens artificially infected with APEC, aiming to provide a theoretical basis and support for the clinical treatment of *E. coli* disease in chickens with MPTA.

## 2 Materials and methods

### 2.1 Ethical Statement

All animal experiments were approved by the animal ethics committee of Hunan Agricultural University, China, approval protocol number: CACAHU 20150418. All procedures involving animals were carried out in strict accordance with the China Laboratory Animal-Welfare Ethical Review Guidelines throughout the experimental process.

### 2.2 Experimental animals

Seventy-five 10-day-old and 350 1-day-old commercial broiler generations of the Lingnan Yellow chicken complete set line (the sire is the yellow-feathered chicken and the dam is the recessive white feathered chicken) (Cao *et al*., 2022) were purchased from Guangdong Zhiwei Agricultural Technology Co. and housed in a strictly sterilized cage system (1.4 m long × 0.7 m wide × 0.4 m high) with controlled temperature and ventilation conditions. The lighting, temperature and relative humidity of the room were adjusted according to the condition of the chickens. The cage was equipped with a feeding trough and a waterer to ensure free feeding and drinking for the test chickens. Antibiotic- and anthelmintic-free basal diets were obtained from Guangdong Nongdao Agricultural Technology Co.

### 2.3 Preparation of MPTA extracts with control drugs

*M. cordata* fruits were collected in Xinning County, Hunan Province, and the samples were preserved at the Key Laboratory of Chinese Veterinary Medicine in Hunan Province. The samples were dried, crushed and extracted by percolation with 0.5% sulfuric acid at 70 °C. The extracts were precipitated with alkali to obtain the crude extracts. The crude extract was then extracted by reflux in 90% ethanol and repeated acid extraction and alkaline precipitation to obtain a brownish MPTA powder with a pungent odor. MPTA Pulvis (MPTA-P) is made from 3.0% MPTA powder mixed with 97.0% corn starch, batch no. 140701. The entire preparation process was carried out in the GMP production plant of Hunan MICOLTA Biological Resources Co., Ltd., and the contents of the main components (PRO ≥ 35%, ALL ≥ 15%) and other trace components in MPTA were pretested by the laboratory (Dong *et al*., 2021, 2022). The doses used in this study were converted from MPTA treatment doses for mice (unpublished). The positive control drug Siwei Chuanxinlian powder (SWCXL-P, consisting of *Andrographis paniculata* (Burm. f.) Nees, *Polygonum hydropiper* L., *Isatis indigotica* Fort. and *Tadehagi triquetrum* (L.) Ohashi) (Tao *et al*., 2020) was supplied by South China Agricultural University Experimental Veterinary Medicine Factory, lot number: 14092801.

### 2.4 Strains and cultures

The test strain *E. coli* O78 of chicken origin (batch number: CVCC1569) was purchased from the China Institute of Veterinary Drugs Control and kept by the School of Veterinary Medicine, South China Agricultural University. Restore glycerol-preserved chicken *E. coli* O78 to room temperature, inoculate aseptically by scratching on normal nutrient medium and incubate for 18-24 h at 37 °C. Individual colonies were picked and inoculated in 5 mL of Luria– Bertani medium and incubated at 37 °C for 16 h. After diluting the bacterial solution into a concentration gradient from 10^−1^ to 10^−10^ according to the plate counting method, 0.1 mL of different concentrations of bacterial solution were taken and spread on agar plates, incubated in a constant temperature incubator at 37 °C for 20 h and then counted. A dish with a colony count of 30-300 was selected as the standard for the determination of the total number of colonies, and each concentration was repeated three times. The concentration of the bacterial solution was adjusted to 2 × 10^9^ CFU/mL.

### 2.5 Development of a pathological model of artificial APEC infection

Seventy-five 10-day-old test chickens were randomly divided into five groups of 15 chickens each, and 0.2 mL, 0.4 mL, 0.6 mL, 0.8 mL and 1.0 mL of *E. coli* broth (concentration of 2×10^9^ CFU/mL) were injected intraperitoneally into each chicken. The dose of bacterial broth that caused 50% to 70% mortality after infection was used as the test dose.

### 2.6 Test animal grouping and treatment

A total of 350 1-day-old test chickens, reared to 10 days of age, and 300 healthy and uniformly weighted (100 ± 5 g) test chickens were randomly divided into 10 groups. Each group of 10 chickens was reared in one cage, and each treatment group contained 3 replicates. Groups 1 to 7 were MPTA-treated (infected, MPTA-P), group 8 was a positive control treatment group (infected, 0.5% SWCXL-P), group 9 was an infected control group (infected, no drug administration), and group 10 was a healthy control group (not infected, no drug administration); the specific dosing regimen is shown in Table 1. Each chicken in groups 1-9 was injected with 0.8 mL of *E. coli* O78 broth, and group 10 was injected with an equal amount of normal saline. Infected and uninfected chickens were kept in separate houses under the same conditions to prevent cross-infection.

**Table 1.**
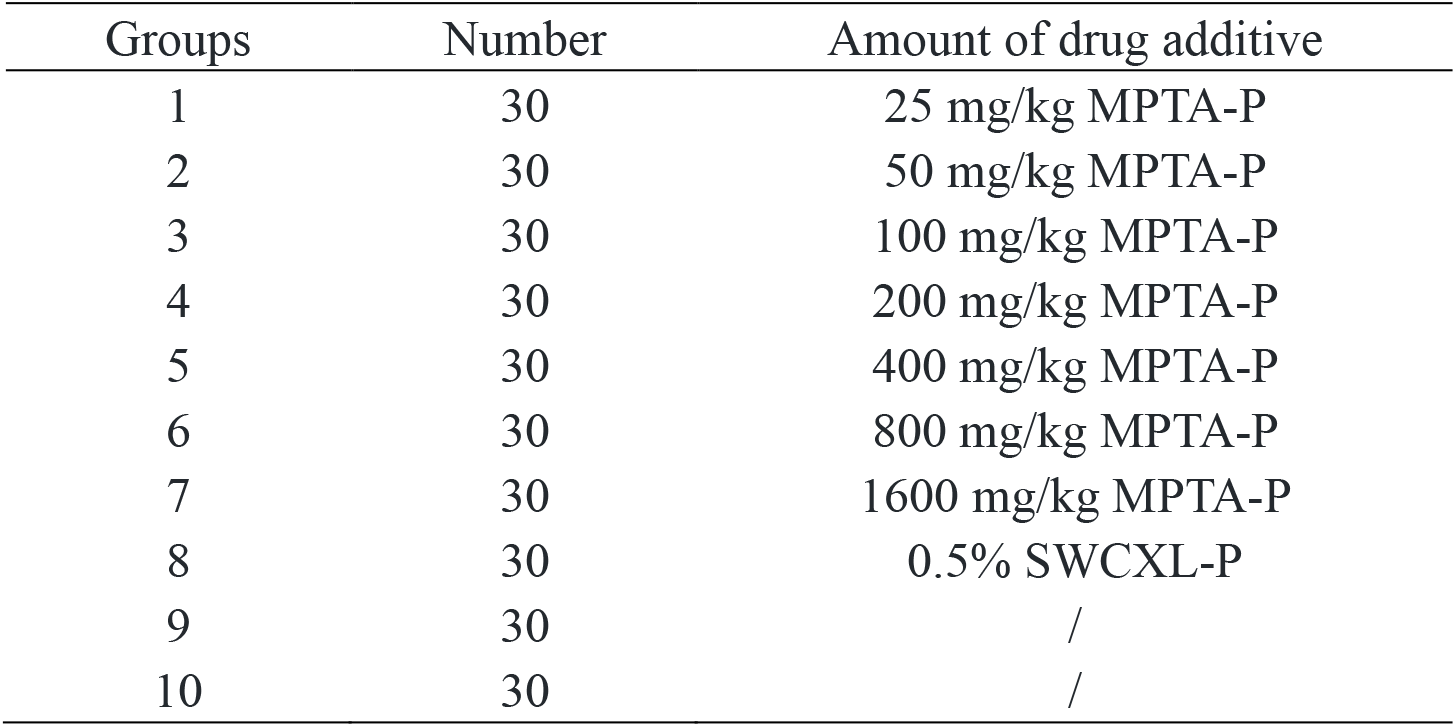
Test groups and treatments.

| Groups | Number | Amount of drug additive |
| --- | --- | --- |
| 1 | 30 | 25 mg/kg MPTA-P |
| 2 | 30 | 50 mg/kg MPTA-P |
| 3 | 30 | 100 mg/kg MPTA-P |
| 4 | 30 | 200 mg/kg MPTA-P |
| 5 | 30 | 400 mg/kg MPTA-P |
| 6 | 30 | 800 mg/kg MPTA-P |
| 7 | 30 | 1600 mg/kg MPTA-P |
| 8 | 30 | 0.5% SWCXL-P |
| 9 | 30 | / |
| 10 | 30 | / |

### 2.7 Clinical signs, mortality and weight

During the test period, the chickens were observed daily for feeding, drinking, diarrhea, mental status and other clinical performance, and the number of sick and dead chickens was recorded. At the end of the test, the surviving chickens were weighed individually, and the average weight of each group was calculated. The surviving but recovered test chickens in each group were then randomly selected for sacrifice and dissected. The effectiveness of therapeutic drugs is judged by the following criteria. Incidence: within 24 hours after artificial inoculation with *E. coli*, test chickens showing signs such as depression, dirty coat, rebellion, bunching, neck shrinkage, drooping wings and loss of appetite are judged to be attacked; Mortality: the pathogenic chickens showed typical symptoms of chicken *E. coli* disease and died, the necropsy showed typical lesion characteristics of chicken *E. coli* disease, and *E. coli* was isolated and cultured from the liver, spleen and heart, which was judged as death by infection; Recovery: The animal is considered to be recovered if its mental state and appetite return to normal after the administration of the drug and if clinical signs such as loose feces no longer appear; Effective: animals that show a significant reduction in clinical signs after administration of the drug and have stopped straining (anal clearing) and are slightly depressed are judged to be effective; Ineffective: animals with clinical signs that have not improved significantly and have even worsened after administration of the drug.

### 2.8 Data statistics and analysis

The test data are expressed as the mean ± standard deviation and were statistically analyzed and processed using SPSS 17.0 software. Body weight was analyzed using one-way ANOVA. The chi-square test was used for the indicators of recovery rate, efficiency rate and mortality rate. P < 0.05 was considered a significant difference.

## 3 Results

### 3.1 Determination of the dose of *E. coli* contamination

The test results showed that after approximately 6-8 h of infection, all chickens showed clinical signs, mainly mental depression, dirty coat, rebellion, pile-up, neck shrinkage, drooping wings, yellow and white droppings, reduced appetite and dead chickens (Fig 1A). Dead chickens were dissected and observed for visceral lesions, which mainly showed enlarged and congested livers, thickened and cloudy air sacs with bleeding spots or bleeding bands, and bleeding spots on the mucosa of the small intestine (Fig 1B-F). Large numbers of typical *E. coli* were isolated from the liver, spleen and heart. The results of the trial showed that clinical signs and bacterial isolation and culture of the test chickens determined that *E. coli* was successfully induced in this trial. During the 7 days of observation, the test results showed that 3, 5, 7, 10 and 13 chickens died in the 0.2 mL to 1.0 mL dose group. Therefore, a dose of 0.8 mL was determined.

**Fig 1.**
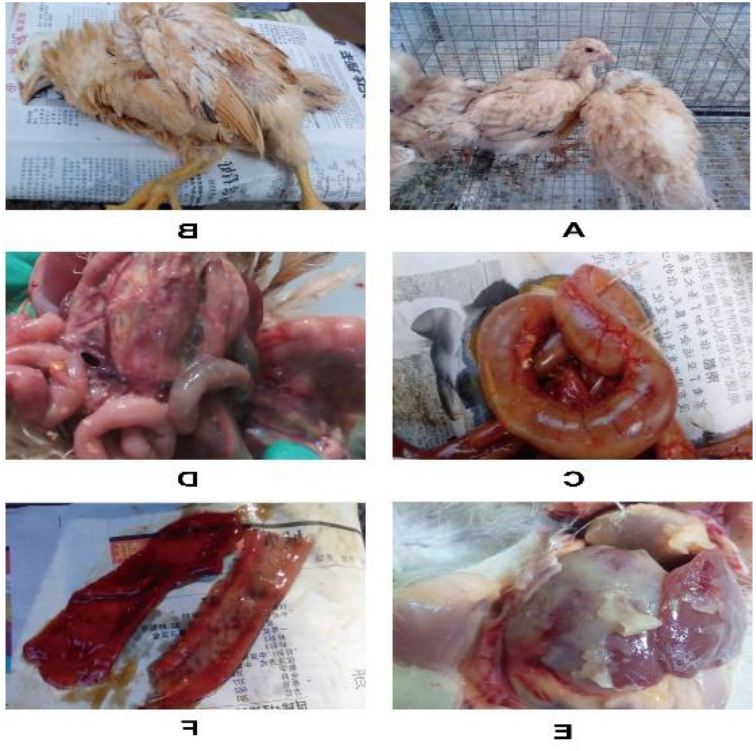
Signs and necropsy of *E. coli* in test chickens. A. Clinical signs in affected chickens; B. Diseased chickens that have died; C. Gas-filled intestines and thinning of the intestinal wall; D. Peritonitis with profuse exudate; E. Perihepatitis, pericarditis and fibrinous exudate; F. Bleeding in the intestinal mucosa

### 3.2 Body weight

All surviving test chickens were weighed at the end of the trial (Table 2). The body weights of test chickens in all infected groups (groups 1 to 9) were significantly lower than those in the control group (group 10) (P<0.05), and the body weights of drug-treated test chickens were significantly higher than those in the infected control group (group 9) (P<0.05). The mean body weight of the test chickens in groups 5 and 6 was significantly higher than that in groups 1 and 2 (P<0.05) and slightly higher than that in groups 3, 4 and 8 (P>0.05). The test results showed that the addition of different doses of MPTA to the diets of infected chickens improved the growth of the test chickens and that groups 5 and 6 improved body weight better than SWCXL-P and other MPTA dose groups.

**Table 2.** Terminal weight of chickens in each test group.

| Groups | Terminal weight |
| --- | --- |
| 1 | 197.47±15.88<sup>e</sup> |
| 2 | 204.53±18.24<sup>de</sup> |
| 3 | 211.63±20.49<sup>cde</sup> |
| 4 | 207.84±19.27<sup>cde</sup> |
| 5 | 222.33±20.99<sup>bc</sup> |
| 6 | 228.38±26.43<sup>b</sup> |
| 7 | 218.45±21.47<sup>bcd</sup> |
| 8 | 206.50±18.09<sup>cde</sup> |
| 9 | 183.00±19.84<sup>f</sup> |
| 10 | 244.78±26.19<sup>a</sup> |

Note: Significant differences between different lowercase letters within the same column (P<0.05).

### 3.3 Clinical signs and mortality

The test results showed that chickens in the nine infected groups showed clinical signs of varying degrees of depression, loss of appetite, loose feathers, drooping wings, closed eyes and shrunken neck, breathing difficulties, standing away from the flock, dysentery, white or yellow watery stools, or in severe cases, lying on the ground within approximately 6 to 8 h after infection. Different doses of the drug were then administered to the affected chickens according to the experimental protocol design.

The highest number of deaths in the test chickens occurred 24 hours after artificial infection, after which the number of deaths gradually decreased. When comparing the time of death of the chickens in each group during the test period, it was found that the chickens in groups 5, 6 and 7 stopped dying after the 6th day of the test, the chickens in groups 1, 3 and 4 stopped dying after the 7th day of the test, the chickens in groups 2 and 8 stopped dying after the 8th day of the test, and the chickens in group 9 stopped dying after the 9th day of the test. By day 10, there were no deaths in any of the test groups. During the trial, nine chickens died in both groups 5 and 6, and the remaining groups also had lower mortality numbers than group 9 (Table 3). The autopsy findings of the infected and dead chickens showed the following characteristics (Fig 2): a peculiar odor that can be smelled when the abdominal cavity is opened; congestion and swelling of the liver with fibrinous exudate on the surface; pericarditis, fibrinous exudate in the epicardium; thickened and cloudy air sacs with exudate of cheese-like material; thinning of the intestinal wall, bleeding of the intestinal mucosa and thinning of the contents. A large number of typical *E. coli* were isolated from the livers, spleens and hearts of dead chickens by bacterial isolation and culture.

**Table 3.**
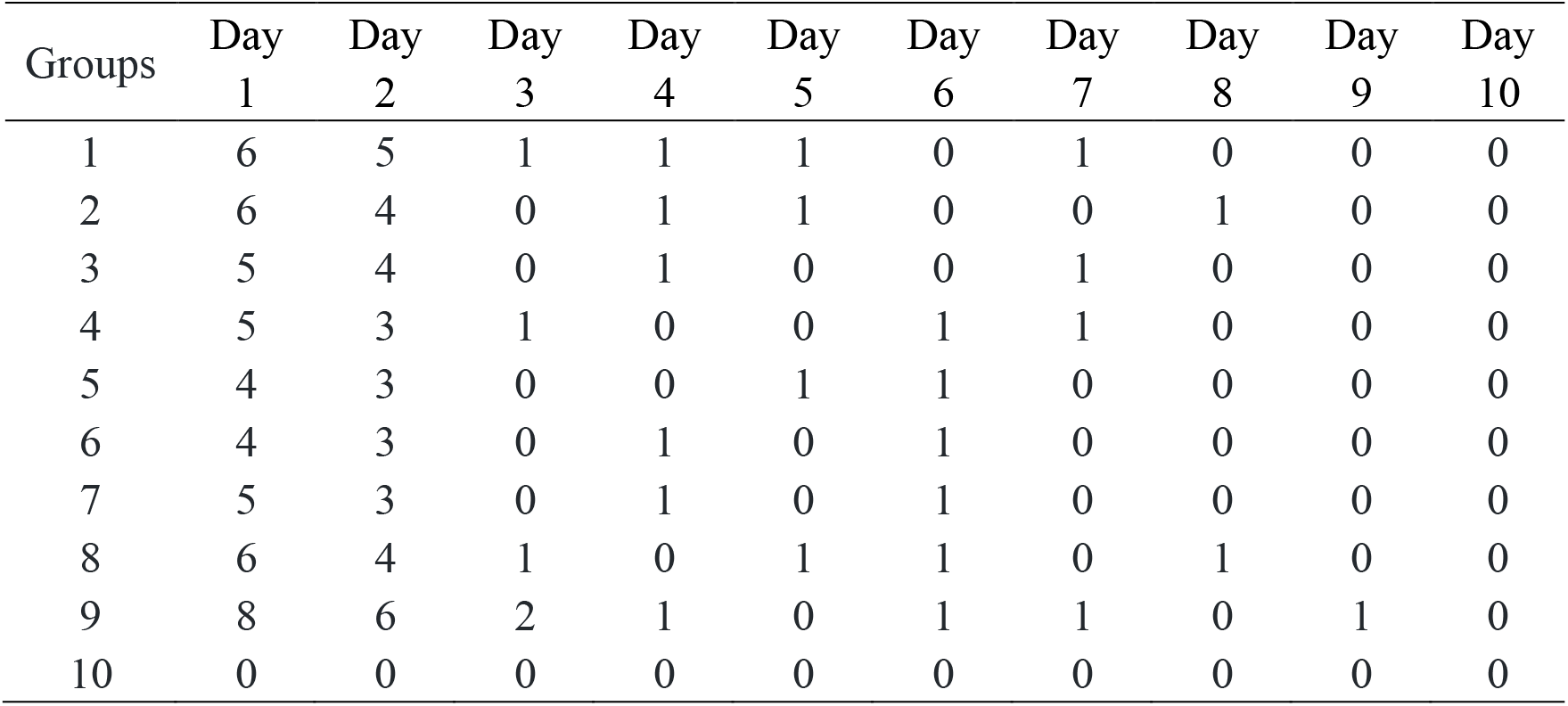
Mortality of test chickens in each group after infection.

| Groups | Day 1 | Day 2 | Day 3 | Day 4 | Day 5 | Day 6 | Day 7 | Day 8 | Day 9 | Day 10 |
| --- | --- | --- | --- | --- | --- | --- | --- | --- | --- | --- |
| 1 | 6 | 5 | 1 | 1 | 1 | 0 | 1 | 0 | 0 | 0 |
| 2 | 6 | 4 | 0 | 1 | 1 | 0 | 0 | 1 | 0 | 0 |
| 3 | 5 | 4 | 0 | 1 | 0 | 0 | 1 | 0 | 0 | 0 |
| 4 | 5 | 3 | 1 | 0 | 0 | 1 | 1 | 0 | 0 | 0 |
| 5 | 4 | 3 | 0 | 0 | 1 | 1 | 0 | 0 | 0 | 0 |
| 6 | 4 | 3 | 0 | 1 | 0 | 1 | 0 | 0 | 0 | 0 |
| 7 | 5 | 3 | 0 | 1 | 0 | 1 | 0 | 0 | 0 | 0 |
| 8 | 6 | 4 | 1 | 0 | 1 | 1 | 0 | 1 | 0 | 0 |
| 9 | 8 | 6 | 2 | 1 | 0 | 1 | 1 | 0 | 1 | 0 |
| 10 | 0 | 0 | 0 | 0 | 0 | 0 | 0 | 0 | 0 | 0 |

**Fig 2.**
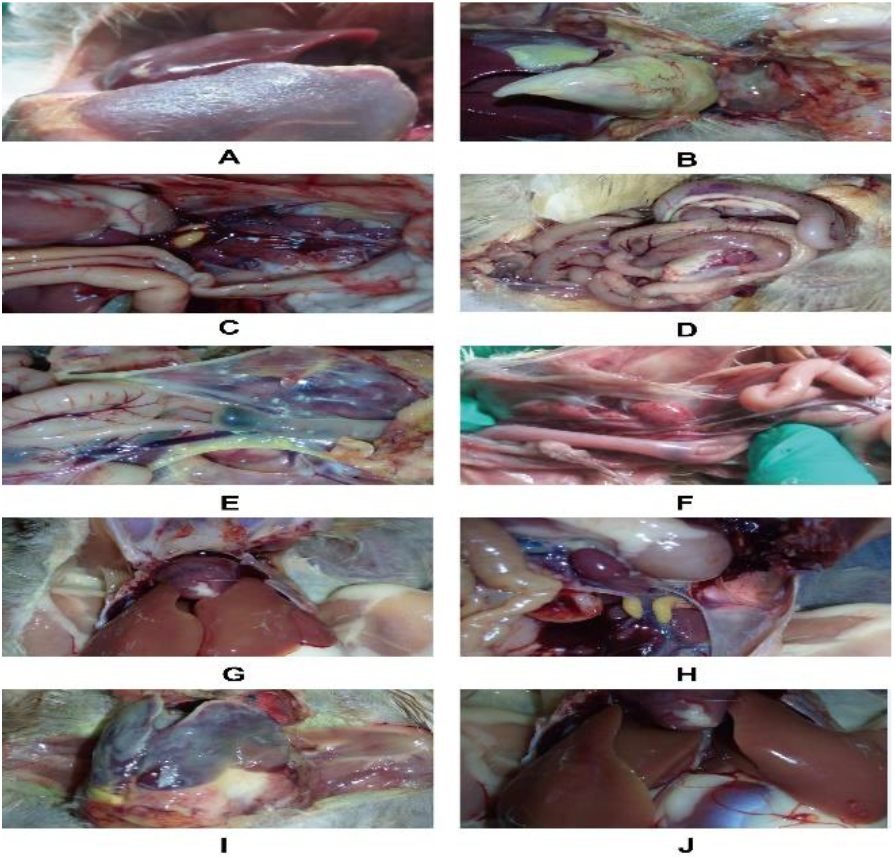
Autopsy of uncured chickens. A to H: the necropsy results for chickens in groups 1-8, in that order; I: the necropsy result of chickens infected with the control group; J: dissection of uninfected chickens.

### 3.4 Efficacy of MPTA in artificially infected chickens with *E. coli* disease

As can be seen from table 4, all groups 1-9 had a 100% incidence rate. The mortality rate in group 9 was 66.7%, which was significantly higher than that in groups 1-8 (p<0.05). The total effective rate results showed that all 8 groups with pharmacological intervention had a higher total effective rate than group 9 (untreated), with groups 2-8 being significantly higher than group 9 (p<0.05). Compared to the positive control group, the total effective rate was higher in groups 3-7 than in group 8, but the difference was not significant (p>0.05).

**Table 4.**
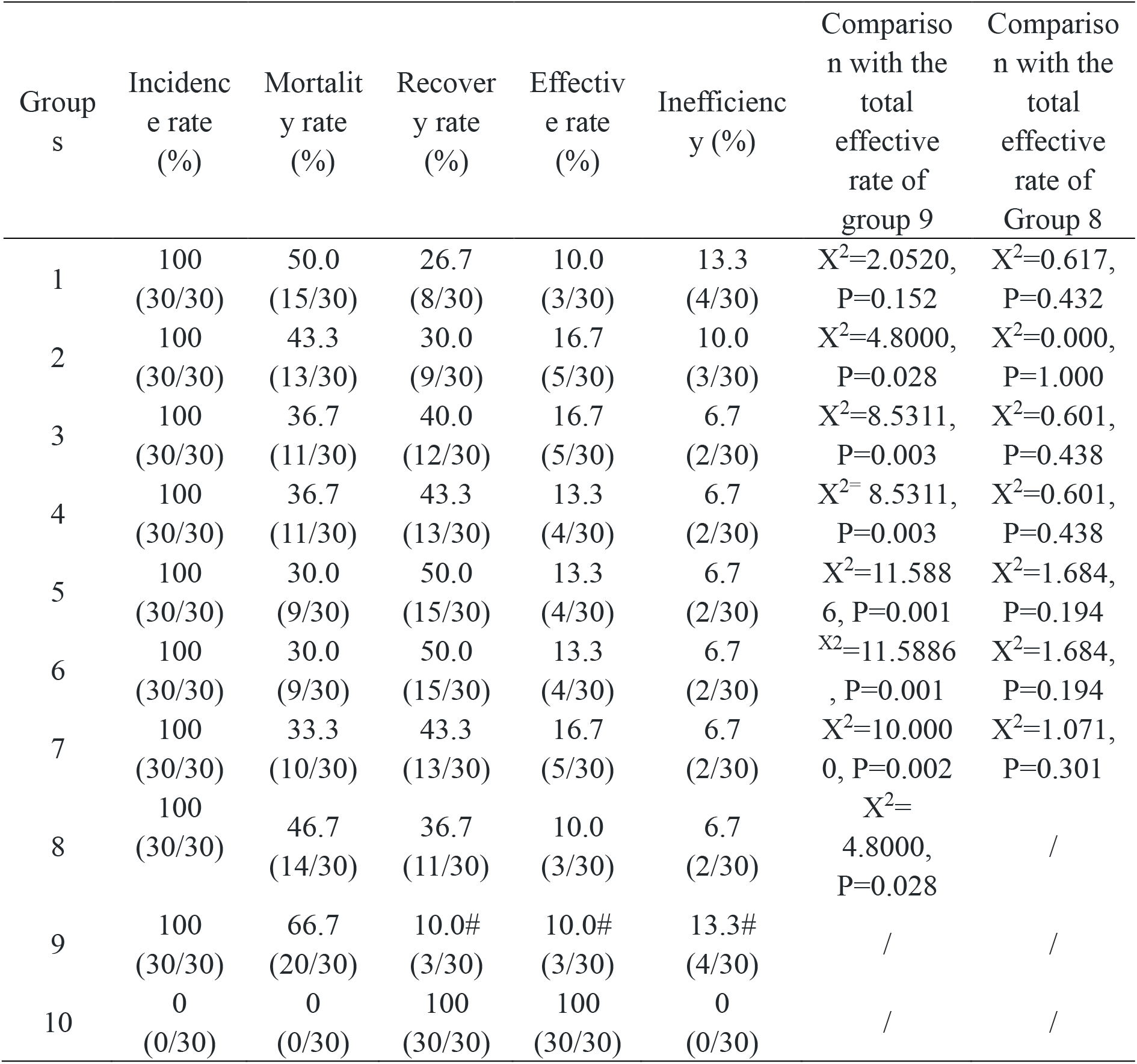
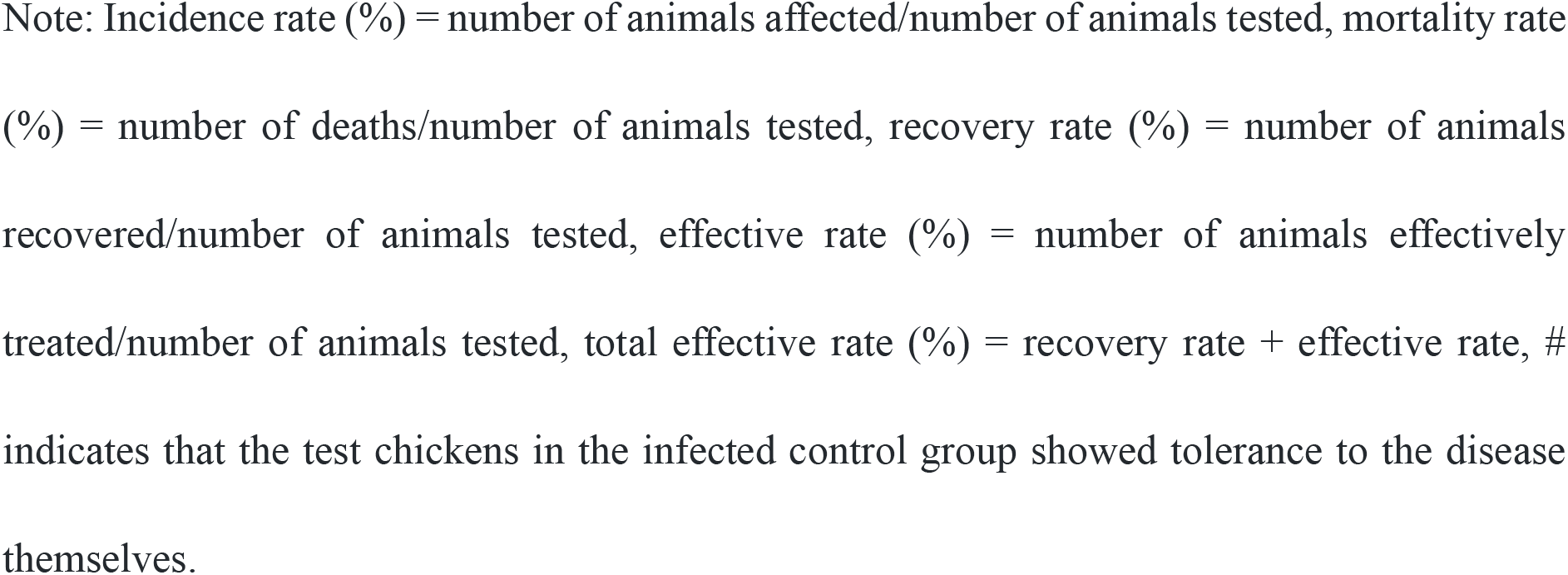
Efficacy of different doses of MPTA and control drug on chicken colibacillosis.

| Group<br>s | Incidence<br>rate<br>(%) | Mortality<br>rate<br>(%) | Recovery<br>rate<br>(%) | Effective<br>rate<br>(%) | Inefficiency<br>(%) | Comparison with the<br>total effective<br>rate of<br>group 9 | Comparison with the<br>total effective<br>rate of<br>Group 8 |
| --- | --- | --- | --- | --- | --- | --- | --- |
| 1 | 100<br>(30/30) | 50.0<br>(15/30) | 26.7<br>(8/30) | 10.0<br>(3/30) | 13.3<br>(4/30) | $X^2=2.0520$,<br>P=0.152 | $X^2=0.617$,<br>P=0.432 |
| 2 | 100<br>(30/30) | 43.3<br>(13/30) | 30.0<br>(9/30) | 16.7<br>(5/30) | 10.0<br>(3/30) | $X^2=4.8000$,<br>P=0.028 | $X^2=0.000$,<br>P=1.000 |
| 3 | 100<br>(30/30) | 36.7<br>(11/30) | 40.0<br>(12/30) | 16.7<br>(5/30) | 6.7<br>(2/30) | $X^2=8.5311$,<br>P=0.003 | $X^2=0.601$,<br>P=0.438 |
| 4 | 100<br>(30/30) | 36.7<br>(11/30) | 43.3<br>(13/30) | 13.3<br>(4/30) | 6.7<br>(2/30) | $X^2=8.5311$,<br>P=0.003 | $X^2=0.601$,<br>P=0.438 |
| 5 | 100<br>(30/30) | 30.0<br>(9/30) | 50.0<br>(15/30) | 13.3<br>(4/30) | 6.7<br>(2/30) | $X^2=11.588$<br>6, P=0.001 | $X^2=1.684$,<br>P=0.194 |
| 6 | 100<br>(30/30) | 30.0<br>(9/30) | 50.0<br>(15/30) | 13.3<br>(4/30) | 6.7<br>(2/30) | $X^2=11.5886$<br>, P=0.001 | $X^2=1.684$,<br>P=0.194 |
| 7 | 100<br>(30/30) | 33.3<br>(10/30) | 43.3<br>(13/30) | 16.7<br>(5/30) | 6.7<br>(2/30) | $X^2=10.000$<br>0, P=0.002 | $X^2=1.071$,<br>P=0.301 |
| 8 | 100<br>(30/30) | 46.7<br>(14/30) | 36.7<br>(11/30) | 10.0<br>(3/30) | 6.7<br>(2/30) | $X^2=4.8000$,<br>P=0.028 | / |
| 9 | 100<br>(30/30) | 66.7<br>(20/30) | 10.0#<br>(3/30) | 10.0#<br>(3/30) | 13.3#<br>(4/30) | / | / |
| 10 | 0<br>(0/30) | 0<br>(0/30) | 100<br>(30/30) | 100<br>(30/30) | 0<br>(0/30) | / | / |

Note: Incidence rate (%) = number of animals affected/number of animals tested, mortality rate (%) = number of deaths/number of animals tested, recovery rate (%) = number of animals recovered/number of animals tested, effective rate (%) = number of animals effectively treated/number of animals tested, total effective rate (%) = recovery rate + effective rate, # indicates that the test chickens in the infected control group showed tolerance to the disease themselves.

### 3.5 Pathologic autopsy

At the end of the trial, three noncured test chickens were randomly selected from each test group for autopsy. Examination revealed minor lesions in the liver, heart, pericardium and small intestine of the infected and medically treated but untreated test chickens (Fig 2A-H). The infected but untreated group 9 test chickens showed more obvious lesions in the liver, heart and small intestine (Fig 2I). The test results showed that the addition of MPTA and SWCXL-P to the diet reduced organ involvement in the test chickens.

## 4 Discussion

The aim of this experiment was to preliminarily evaluate the effectiveness and dosage of MPTA in the control of APEC infections in chickens. After successful artificial infection, all MPTA groups showed varying degrees of reduction in the in vivo effects of *E. coli* disease, better weight recovery, lower mortality and shorter duration of clinical signs compared to infected controls. These findings suggest that the ameliorative effect of MPTA on *E. coli* may reduce the suffering of affected chickens, improve animal welfare and reduce the economic losses associated with the disease.

There is sufficient evidence that invasive *E. coli* infections can cause the body to enter a sustained state of oxidative stress and produce widespread inflammatory damage (Baronetti *et al*., 2013; Croxen and Finlay, 2010; Farr and Kogoma, 1991; Fu *et al*., 2022). The protective effect of MPTA against *E. coli* disease may be due to immunomodulation and oxidative stress protection found in previous studies and in the literature (Alam *et al*., 2019; Dong *et al*., 2022; Nigdelioglu Dolanbay, Kocanci, and Aslim, 2021; Zhang *et al*., 2019). Compared to the infected control group, chickens in both the MPTA-supplemented and SWCXL-P groups showed significant improvements in body weight suppression, with the greatest recovery occurring at the 400 and 800 mg/kg supplementation doses. Improved growth rates due to MPTA supplementation may be attributed to the regulation of intestinal flora, inflammation inhibition and free radical scavenging (Liu *et al*., 2022).

Studies have shown that MPTA supplementation can increase albumin and globulin concentrations in laying hens, thereby activating the complement system and establishing an adaptive immune response to achieve at least partial pathogen clearance and defense against infection (Basta, 2008; van Dijk *et al*., 2018; Miletic and Frank, 1995). Over the course of the disease, supplementation with 400-1600 mg/kg of MPTA-P stopped the death of the affected chickens by day 7 of treatment, accelerating recovery from the disease. The results of the gross pathological profile showed that MPTA extract significantly reduced inflammation and damage to various organs caused by *E. coli* disease. It was concluded that dietary supplementation with 400 mg/kg MPTA-P for 7 consecutive days significantly reduced the clinical course and signs of *E. coli*-infected broilers and provided protection against *E. coli* infection and organ damage.

## Reference

Alam MB, Ju M-K, Kwon Y-G, et al., 2019. Protopine attenuates inflammation stimulated by carrageenan and LPS via the MAPK/NF-κB pathway. Food Chem Toxicol Int J Publ Br Ind Biol Res Assoc 131:110583.

Bae DS, Kim YH, Pan C-H, et al., 2012. Protopine reduces the inflammatory activity of lipopolysaccharide-stimulated murine macrophages. BMB Rep 45:108–113.

Baronetti JL, Villegas NA, Aiassa V, et al., 2013. Hemolysin from Escherichia coli induces oxidative stress in blood. Toxicon Off J Int Soc Toxinology 70:15–20.

Basta M, 2008. Ambivalent effect of immunoglobulins on the complement system: activation versus inhibition. Mol Immunol 45:4073–4079.

Cao N, Yang S, Hu C, et al., 2022. Effects of Polysaccharide of Atractylodes macrocephala Koidz on Growth Performance, Slaughter Performance and Immune Organ Development of Lingnan Yellow Chickens. Chin J Anim Nutr 34:205–214.

Christensen H, Bachmeier J and Bisgaard M, 2021. New strategies to prevent and control avian pathogenic Escherichia coli (APEC). Avian Pathol J WVPA 50:370–381.

Croxen MA and Finlay BB, 2010. Molecular mechanisms of Escherichia coli pathogenicity. Nat Rev Microbiol 8:26–38.

Dakheel MM, Alkandari F a. H, Mueller-Harvey I, et al., 2020. Antimicrobial in vitro activities of condensed tannin extracts on avian pathogenic Escherichia coli. Lett Appl Microbiol 70:165–172.

Dho-Moulin M and Fairbrother JM, 1999. Avian pathogenic Escherichia coli (APEC). Vet Res 30:299–316.

van Dijk A, Hedegaard CJ, Haagsman HP, et al., 2018. The potential for immunoglobulins and host defense peptides (HDPs) to reduce the use of antibiotics in animal production. Vet Res 49:68.

Dong Z, Liu M, Zhong X, et al., 2021. Identification of the Impurities in Bopu Powder® and Sangrovit® by LC-MS Combined with a Screening Method. Molecules 26:3851.

Dong Z, Wang Y, Tang Z, et al., 2022. Exploring the Anti-inflammatory Effects of Protopine Total Alkaloids of Macleaya Cordata (Willd.) R. Br. Front Vet Sci 9.

Dos Anjos Adur M, Châtre P, Métayer V, et al., 2022. Escherichia coli ST224 and IncF/blaCTX-M-55 plasmids drive resistance to extended-spectrum cephalosporins in poultry flocks in Parana, Brazil. Int J Food Microbiol 380:109885.

Dube N and Mbanga J, 2018. Molecular characterization and antibiotic resistance patterns of avian fecal Escherichia coli from turkeys, geese, and ducks. Vet World 11:859–867.

Eid S, Tolba HMN, Hamed RI, et al., 2022. Bacteriophage therapy as an alternative biocontrol against emerging multidrug resistant E. coli in broilers. Saudi J Biol Sci 29:3380–3389.

Farr SB and Kogoma T, 1991. Oxidative stress responses in Escherichia coli and Salmonella typhimurium. Microbiol Rev 55:561–585.

Fu Q, Lin Q, Chen D, et al., 2022. β-defensin 118 attenuates inflammation and injury of intestinal epithelial cells upon enterotoxigenic Escherichia coli challenge. BMC Vet Res 18:142.

Ghunaim H, Abu-Madi MA and Kariyawasam S, 2014. Advances in vaccination against avian pathogenic Escherichia coli respiratory disease: potentials and limitations. Vet Microbiol 172:13–22.

Guo X, Zhang L-Y, Wu S-C, et al., 2014. Andrographolide interferes quorum sensing to reduce cell damage caused by avian pathogenic Escherichia coli. Vet Microbiol 174:496–503.

Helmy YA, Deblais L, Kassem II, et al., 2018. Novel small molecule modulators of quorum sensing in avian pathogenic Escherichia coli (APEC). Virulence 9:1640–1657.

Helmy YA, Kathayat D, Deblais L, et al., 2022. Evaluation of Novel Quorum Sensing Inhibitors Targeting AutoInducer 2 (AI-2) for the Control of Avian Pathogenic Escherichia coli Infections in Chickens. Microbiol Spectr 10:e0028622.

Johnson TJ, Miller EA, Flores-Figueroa C, et al., 2022. Refining the definition of the avian pathogenic Escherichia coli (APEC) pathotype through inclusion of high-risk clonal groups. Poult Sci 101:102009.

Jørgensen F, McLauchlin J, Verlander NQ, et al., 2022. Levels and genotypes of Salmonella and levels of Escherichia coli in frozen ready-to-cook chicken and turkey products in England tested in 2020 in relation to an outbreak of S. Enteritidis. Int J Food Microbiol 369:109609.

Ke W, Lin X, Yu Z, et al., 2017. Molluskicidal activity and physiological toxicity of Macleaya cordata alkaloids components on snail Oncomelania hupensis. Pestic Biochem Physiol 143:111–115.

Ke W, Tu C, Cao D, et al., 2019. Molluskicidal activity and physiological toxicity of quaternary benzo[c]phenanthridine alkaloids (QBAs) from Macleaya cordata fruits on Oncomelania hupensis. PLoS Negl Trop Dis 13:e0007740.

Koutsianos D, Athanasiou LV, Mossialos D, et al., 2022. Investigation of Serotype Prevalence of Escherichia coli Strains Isolated from Layer Poultry in Greece and Interactions with Other Infectious Agents. Vet Sci 9:152.

Kulnanan P, Chuprom J, Thomrongsuwannakij T, et al., 2021. Antibacterial, antibiofilm, and anti-adhesion activities of Piper betle leaf extract against Avian pathogenic Escherichia coli. Arch Microbiol 204:49.

Liang W, Li H, Zhou H, et al., 2021. Effects of Taraxacum and Astragalus extracts combined with probiotic Bacillus subtilis and Lactobacillus on Escherichia coli-infected broiler chickens. Poult Sci 100:101007.

Liu H, Lin Q, Liu X, et al., 2022. Effects of Dietary Bopu Powder Supplementation on Serum Antioxidant Capacity, Egg Quality, and Intestinal Microbiota of Laying Hens. Front Physiol 13:902784.

Lutful Kabir SM, 2010. Avian colibacillosis and salmonellosis: a closer look at epidemiology, pathogenesis, diagnosis, control and public health concerns. Int J Environ Res Public Health 7:89–114.

Meena PR, Yadav P, Hemlata H, et al., 2021. Poultry-origin extraintestinal Escherichia coli strains carrying the traits associated with urinary tract infection, sepsis, meningitis and avian colibacillosis in India. J Appl Microbiol 130:2087–2101.

Miletic VD and Frank MM, 1995. Complement-immunoglobulin interactions. Curr Opin Immunol 7:41–47.

Muñoz O, Christen P, Cretton S, et al., 2011. Chemical study and anti-inflammatory, analgesic and antioxidant activities of the leaves of Aristotelia chilensis (Mol.) Stuntz, Elaeocarpaceae. J Pharm Pharmacol 63:849–859.

Nigdelioglu Dolanbay S, Kocanci FG and Aslim B, 2021. Neuroprotective effects of allocryptopine-rich alkaloid extracts against oxidative stress-induced neuronal damage. Biomed Pharmacother Biomedecine Pharmacother 140:111690.

Oliveira JM, Cardoso MF, Moreira FA, et al., 2022. Phenotypic antimicrobial resistance (AMR) of avian pathogenic Escherichia coli (APEC) from broiler breeder flocks between 2009 and 2018. Avian Pathol J WVPA 51:388–394.

Rabiu AG, Falodun OI, Fagade OE, et al., 2022. Potentially pathogenic Escherichia coli from household water in peri-urban Ibadan, Nigeria. J Water Health 20:1137–1149.

Sarowska J, Olszak T, Jama-Kmiecik A, et al., 2022. Comparative Characteristics and Pathogenic Potential of Escherichia coli Isolates Originating from Poultry Farms, Retail Meat, and Human Urinary Tract Infection. Life Basel Switz 12:845.

Tao S, Wang Y, Liao C, et al., 2020. Bacteriostatic effects of Siweichuanxinlian powder and its components on Salmonella in vitro. J Tradit Chin Vet Med 39:64–67.

Zhang B, Zeng M, Li M, et al., 2019. Protopine Protects Mice against LPS-Induced Acute Kidney Injury by Inhibiting Apoptosis and Inflammation via the TLR4 Signaling Pathway. Mol Basel Switz 25:E15.

Zhang Q, Zhang Z, Zhou S, et al., 2021. Macleaya cordata extract, an antibiotic alternative, does not contribute to antibiotic resistance gene dissemination. J Hazard Mater 412:125272.

Zhong X, Shi Y, Chen J, et al., 2014. Polyphenol extracts from Punica granatum and Terminalia chebula are anti-inflammatory and increase the survival rate of chickens challenged with Escherichia coli. Biol Pharm Bull 37:1575–1582.

